# The urinary microbiome associated with bladder cancer

**DOI:** 10.1101/224667

**Authors:** Viljemka Bučević Popović, Marijan Šitum, Cheryl-Emiliane T. Chow, Luisa S. Chan, Blanka Roje, Janoš Terzić

## Abstract

Recent findings suggest that microorganisms inhabiting the human body can influence the development of cancer, but the role of microorganisms in bladder cancer pathogenesis has not been explored yet. The aim of this study was to characterize and compare the urinary microbiome of bladder cancer patients with those of healthy controls. Bacterial communities present in urine specimens collected from male patients diagnosed with primary or recurrent, non-muscle invasive bladder cancers, and from healthy, age-matched individuals were analysed using 16S Illumina MiSeq sequencing. Our result show that the most abundant phylum in both groups was *Firmicutes*, followed by *Actinobacteria, Bacteroidetes* and *Proteobacteria*. While microbial diversity and overall microbiome composition were not significantly different between bladder cancer and healthy samples, we identified specific operational taxonomic units (OTUs) that were significantly more abundant (p < 0.05) in either type of samples. Among those that were significantly enriched in the bladder cancer group, we identified an OTU belonging to genus *Fusobacterium*, a possible protumorigenic pathogen. Three OTUs more abundant in healthy urines were from genera *Veillonella, Streptococcus* and *Corynebacterium*. Detected microbiome changes suggest that microbiome may be a factor in bladder cancer pathology, and the clinical implications of reported results remain to be explored.

## Introduction

Bladder cancer is the ninth most frequent malignant disease, with more than 160,000 deaths per year reported globally. The risk of developing the disease increases with age, and it is diagnosed three times more often in men than in women. Because majority of new cases are found in people above 65 years of age due to increased life expectancy, it is anticipated that the number of affected individuals will surge in the future^1^.

Apart from environmental and genetic risk factors, researchers have become increasingly aware that microbes inhabiting the human body play an important role for maintenance of health and the development of disease. Microbiome studies, fuelled by the availability of high-throughput DNA-based techniques, have shown that perturbation in the microbiome is associated with a number of human diseases. The vast majority of these studies was performed on the gut, the body niche where most of commensal microorganisms reside, and associations were found between microbiome and diseases such as inflammatory bowel disease, multiple sclerosis, type 1 and 2 diabetes, allergies, asthma, autism, as well as cancer^2^.

The link between cancer and specific microbial agents is well known and it is estimated that microorganisms contributes to up to 20% of human malignancies^3^. The most prominent examples are *Helicobacter pylori* implicated in the development of gastric cancer, and high-risk types of human papillomavirus in cervical cancer^4^. The interaction of microorganisms and their hosts is extremely complex, and a multitude of molecular mechanisms may be envisioned by which they influence oncogenesis, tumour progression and response to anticancer therapy^3,5–6^. Bacteria can directly damage host DNA via genotoxins, such as colibactin produced by some *E. coli* strains, or indirectly by generating reactive oxidative species. Some pathogenic microorganisms manipulate host signalling pathways, exemplified by Wnt/β-catenin pathway which is altered to support cell proliferation in many types of cancers. Microbiome can also induce chronic inflammation providing a background for tumour development or elicit immunosuppressive responses that may subvert cancer immunosurvellience. Finally, bacterial metabolism of host derived metabolites, food components or xenobiotics may result in harmful compounds that may promote tumorigenesis even at distant body sites^3,5^.

Traditionally, bladder epithelium and urine have been considered sterile in healthy individuals. This assumption was based primarily on microbiological urine cultures, best suited for detecting aerobic, fast-growing uropathogens. The evidence has accumulated during the last five years, that urinary tract also harbours distinct commensal microorganisms^8^. The urinary microbiome reported for healthy people varies considerably due to use of different analytical and urine collection methods. While female urinary microbiome is much better characterized than male, it can be concluded that urinary microbiome is marked by sex- and age-related differences as well as significant inter-individual variability^9,10^. Studies have explored the changes in urinary microbiome in states such as type 2 diabetes mellitus^11^, overactive bladder^12,13^, urinary incontinence ^14–16^, interstitial cystitis^17^, neuropathic bladder^18,19^, sexually transmitted infections^20^ or chronic prostatitis/chronic pelvic pain syndrome^21,22^. The urinary microbiome in urothelial bladder cancer has not been investigated, apart from the pilot study by Xu *et al*.^23^ that reported enrichment of *Streptococcus* sp. in some of the cancer patients. Our study characterized the urinary microbiome of bladder cancer patients and compared it with that of healthy controls to gain insight into the microbiome’s possible role in bladder cancer.

## Results

### Participant characteristics and sequencing data summary

A total of 36 subjects were included in the study. However, 12 samples failed to provide sufficient DNA for sequencing and one sample did not meet sequencing quality criteria due to low sequencing depth. Supplementary Table S1 displays characteristics of remaining 23 subjects (12 bladder cancer patients and 11 healthy controls).

Sequencing of urine samples plus extraction control resulted with a total of 22,341,934 raw sequences. These were merged into 9,977,955 paired sequences, with an average read length of 252 base pairs. Filtering for sequence quality and OTU prevalence (min. 10% of samples) reduced the number of sequences to 9,713,510, assigned to 348 OTUs. One of the bladder cancer samples with less than 50,000 reads was excluded from further analysis. Rarefaction curves show that the remaining 23 samples were sequenced to a sufficient depth such that a complete microbiome profile was likely captured for most samples (Supplementary Fig. S1). Classification to the genus level was possible for 95% of sequence reads. A total of 10 bacterial phyla, 19 classes, 26 orders, 61 families and 107 genera were identified. Only two OTUs were detected in the extraction control, one belonged to genus *Bacteroides* and the other was a chloroplast from the order *Streptophyta*.

### Microbiome diversity and composition of bladder cancer and healthy urine samples

Both metrics used to assess differences in microbial alpha diversity (richness and Shannon Index) were statistically similar between cancerous and healthy samples (Fig. 1). The average number of observed OTUs found within a sample was 182 for the bladder cancer group and 184 for healthy controls (Fig. 1a).

**Figure 1.**
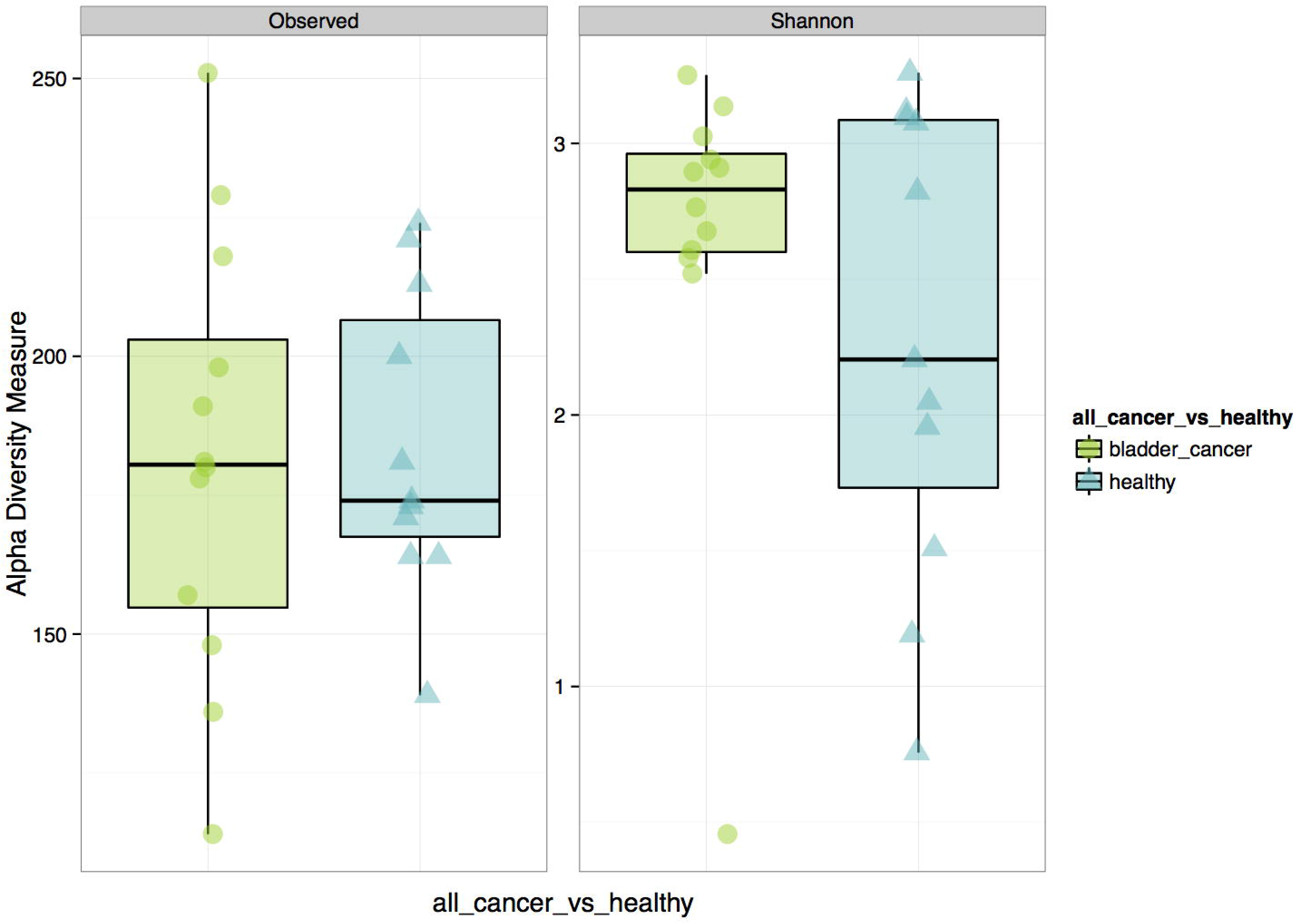
Microbial alpha diversity of urine samples. Bladder cancer patients (green circles); Healthy males (blue triangles): **(a)** Observed number of OTUs, **(b)** Shannon Index. Both alpha diversity metrics were statistically similar between cancer and healthy samples.

The urinary microbiome of bladder cancer patients and healthy controls at the phylum and the family level is shown in Fig. 2. The most abundant phyla included *Firmicutes, Actinobacteria, Bacteroidetes* and *Proteobacteria*. A prominent feature of urinary microbiome evidenced from these results is that there is a high degree of inter-individual variability in community composition among study participants in both bladder cancer and healthy subgroups. The microbial composition of the sample collected from one of the bladder cancer patients (AK15_4004 in Fig. 2) was inconsistent with other urine samples; it was dominated by the family *Enterobacteriaceae* with relative abundance of 91% and it was excluded from community structure analysis.

**Figure 2.**
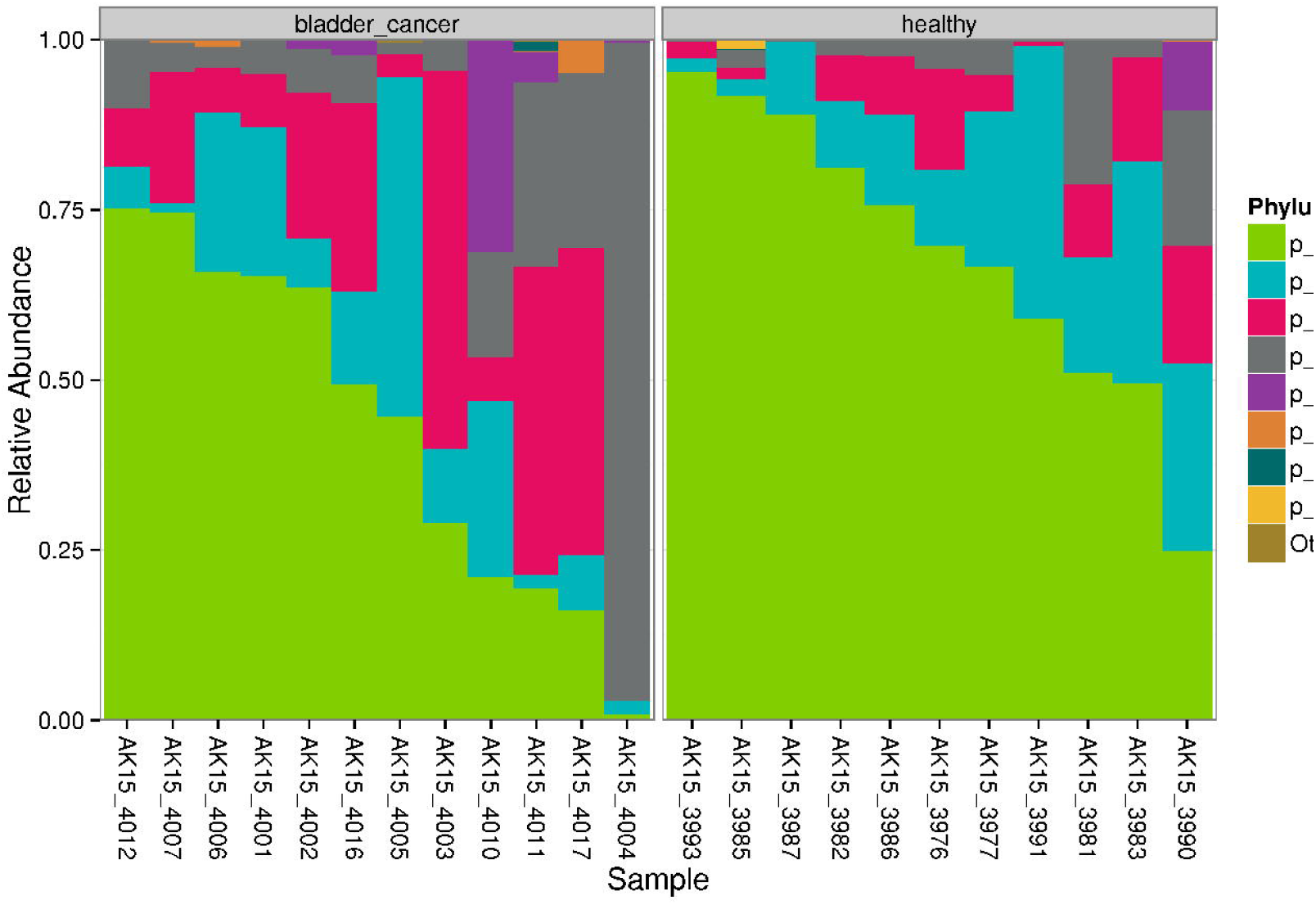
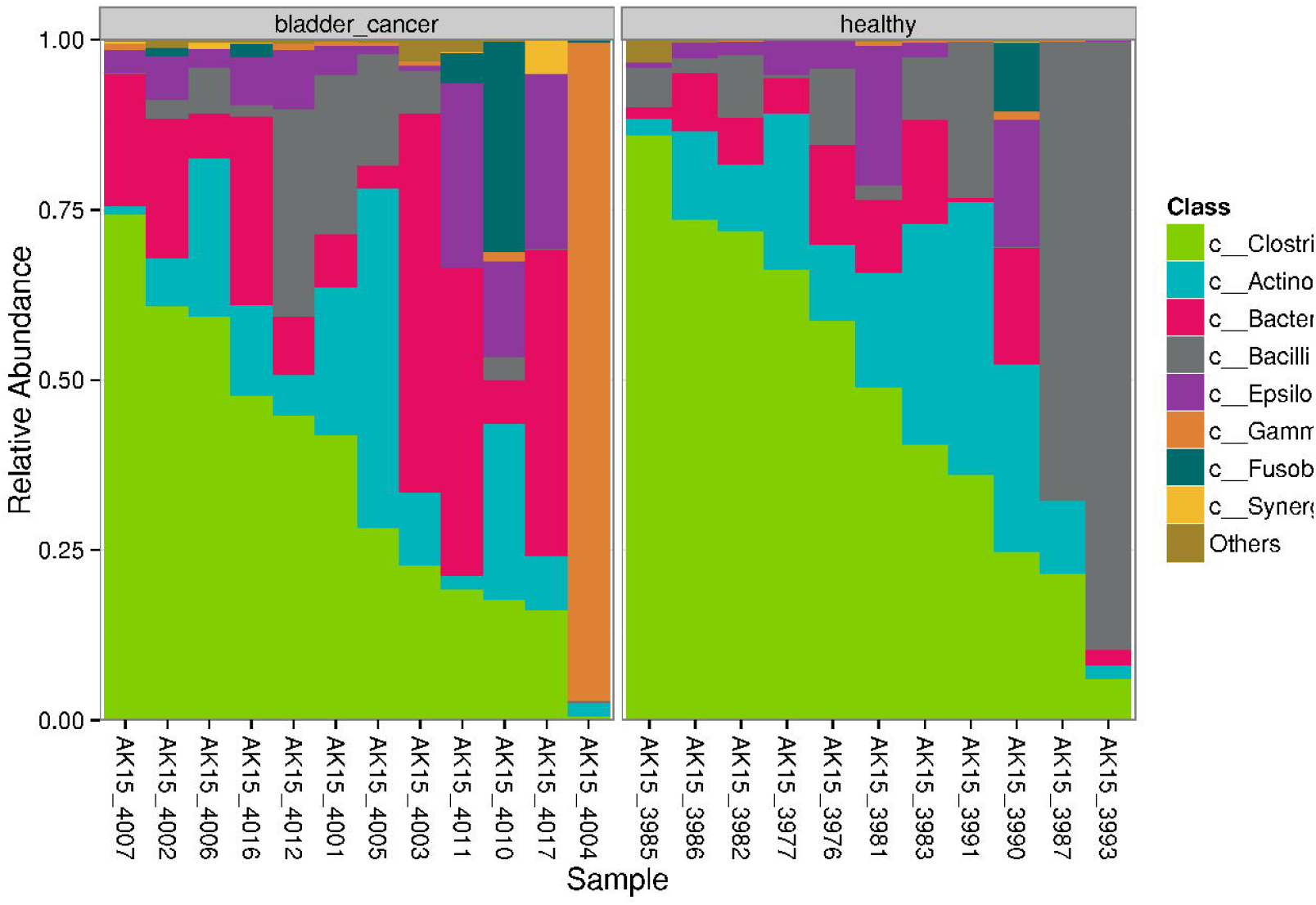
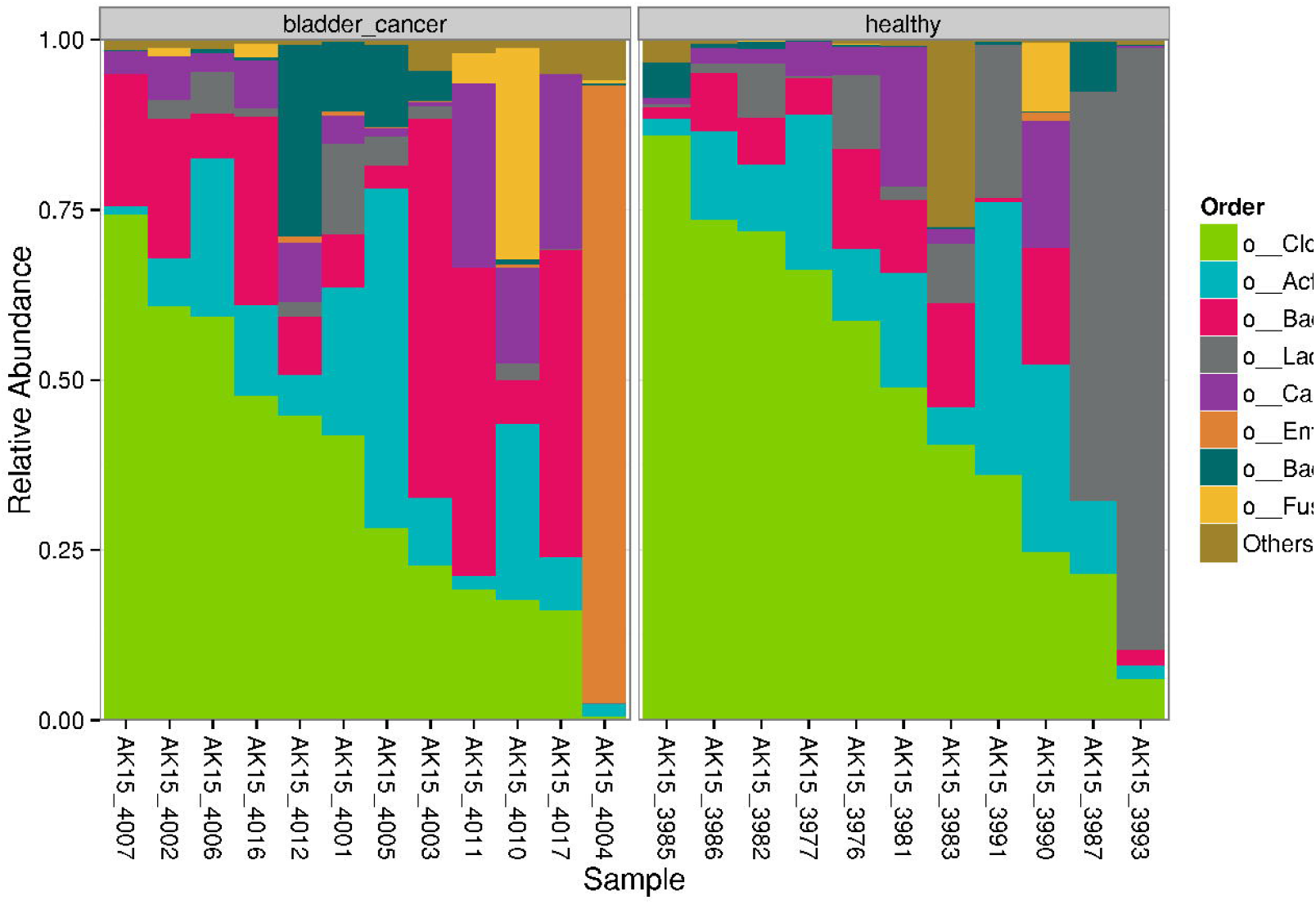
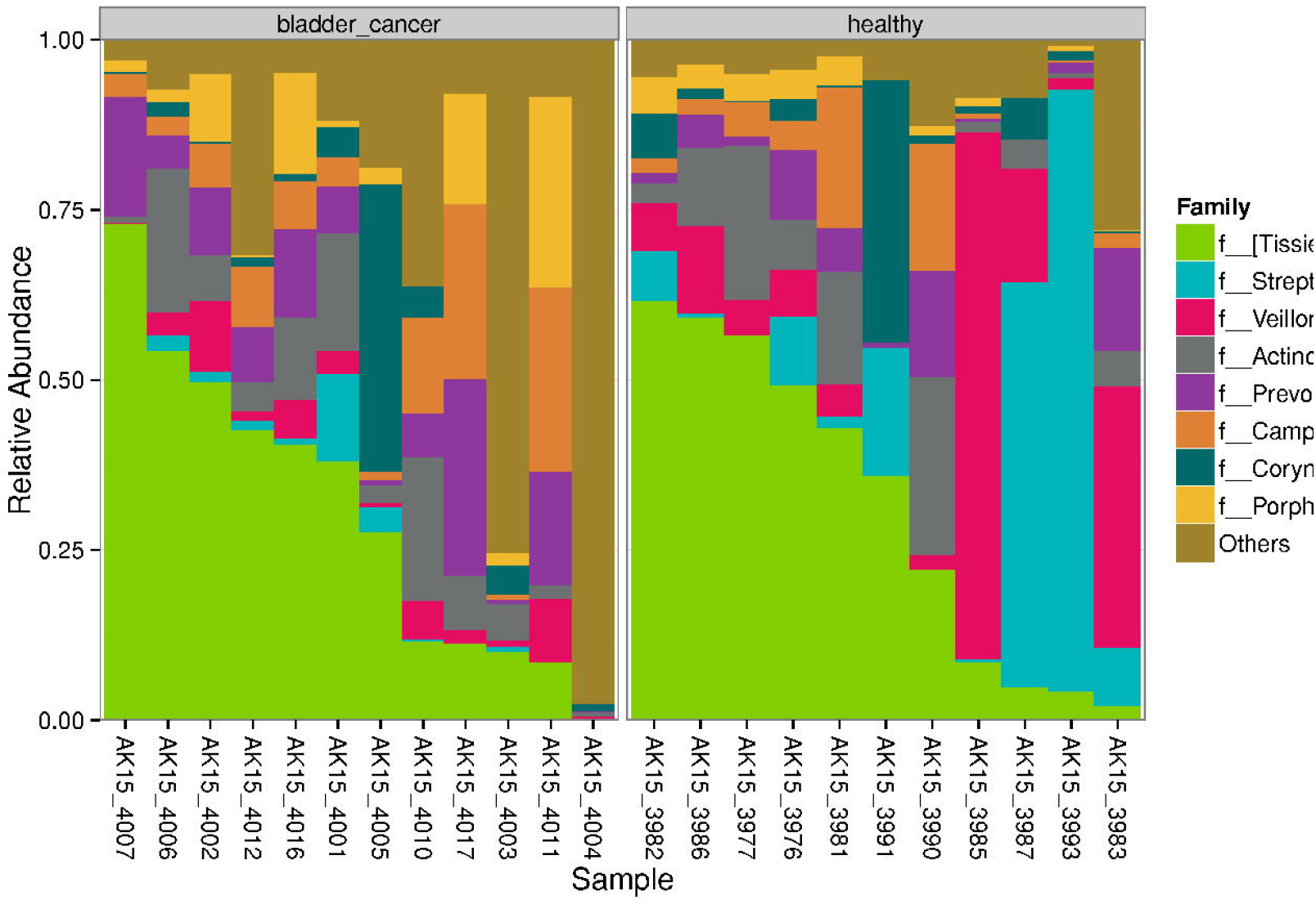
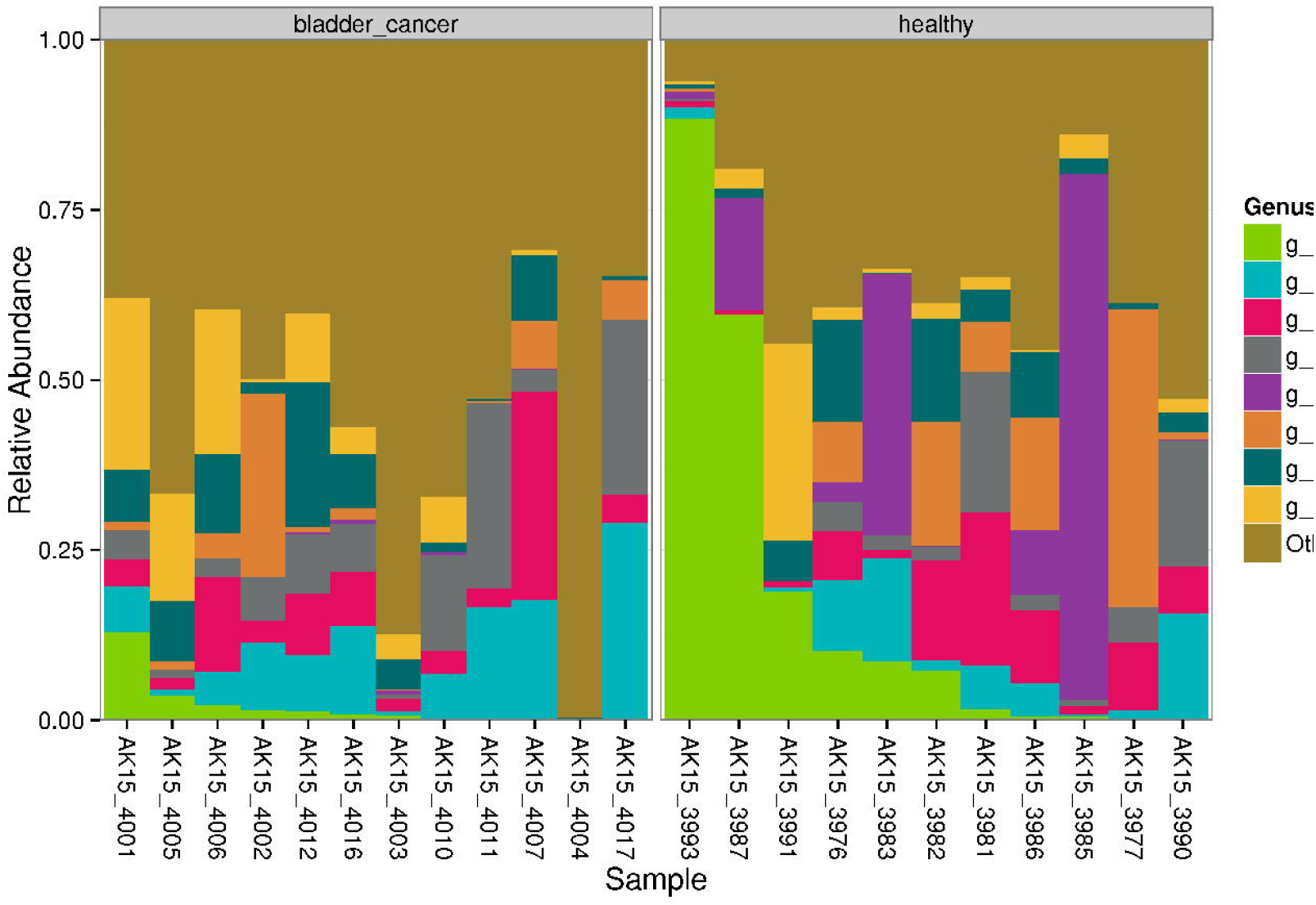
Urinary microbiota of male bladder cancer patients and healthy controls. Most abundant taxa are shown at phylum **(a)**, class **(b)**, order **(c)**, family **(d)** and genus **(e)** level.

No differences in the overall urinary microbiome of bladder cancer urine samples compared to healthy samples was observed as bladder cancer urine samples did not cluster in the PCoA (Fig. 3). A PERMANOVA analysis was also performed to determine if there is a significant association between microbiome composition and other tested variables such as malignancy, cancer type or patient age. Among those, variations in the urine microbiome were significantly associated only with age across all samples (p = 0.008).

**Figure 3.**
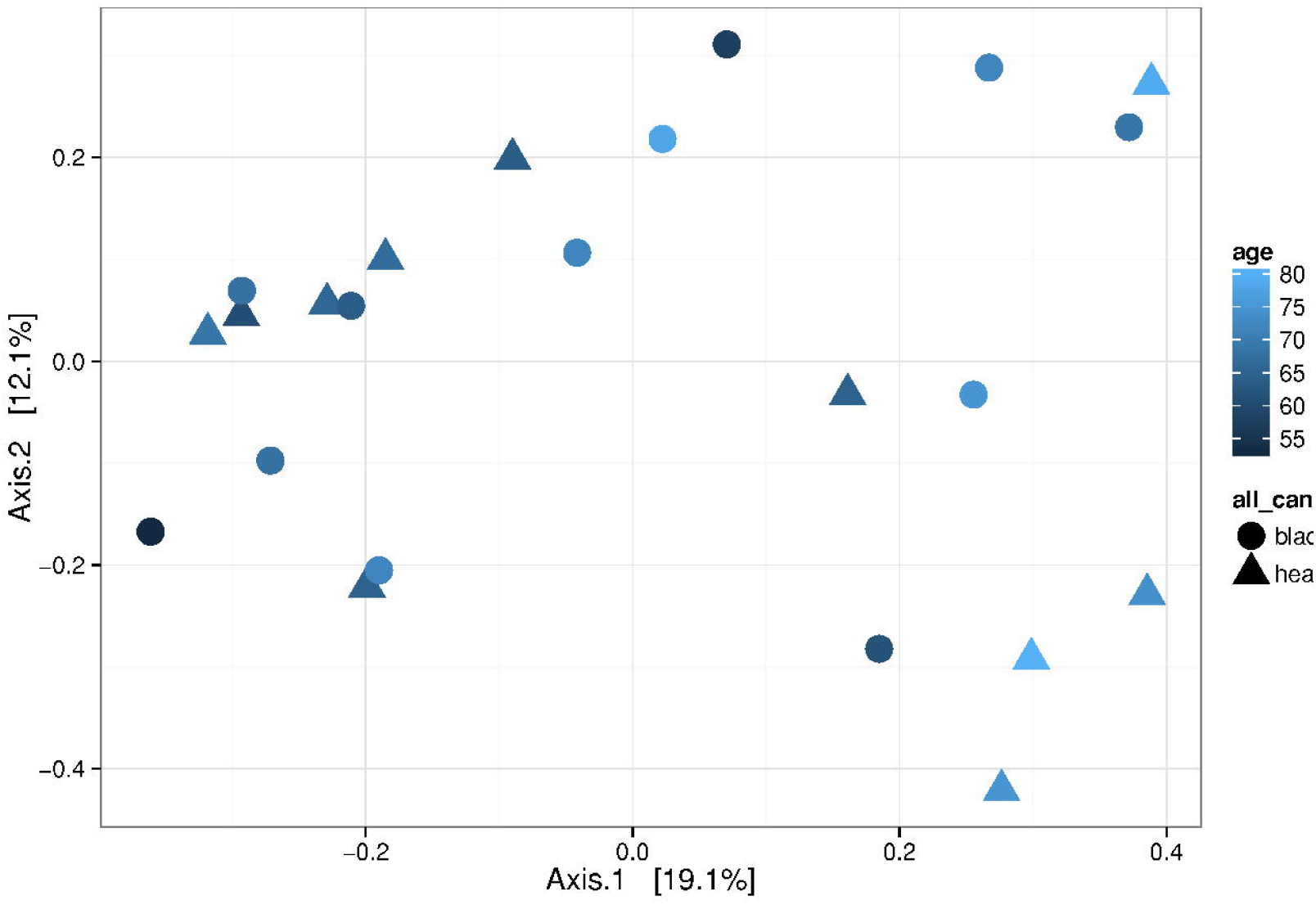
Microbial beta diversity. Dimensional reduction of the Bray-Curtis distance between microbiome samples, using PCoA ordination method, for bladder cancer urines (dots) and healthy controls (triangles). Data points are coloured according to age in years. Permutational analysis of variance shows that variations in the urine microbiome were significantly associated with age (p = 0.008). Samples do not cluster according to their cancer/healthy status.

### Community structure reveals differently abundant OTUs in bladder cancer and healthy urine

While there was no significant difference in microbiome composition in terms of overall diversity or composition at the phylum or family level, specific OTUs were identified that exhibited significant differences (p < 0.05) in abundance between cancer and healthy samples (Fig. 4). Eight OTUs were enriched in urine of bladder cancer patients including genera *Fusobacterium, Actinobaculum, Facklamia, Campylobacter*. Three OTUs identified at strain level were *Campylobacter hominis, Actinobaculum massiliense*, and *Jonquetella antropi* (94otu40402). Five OTUs were enriched in healthy samples from the genera *Veillonella, Streptococcus, Corynebacterium*; these OTUs were further identified as *Veillonella dispar* at species, and *Streptococcus cristatus, Corynebacterium appendicis* and *Corynebacterium sp*. at strain level.

**Figure 4.**
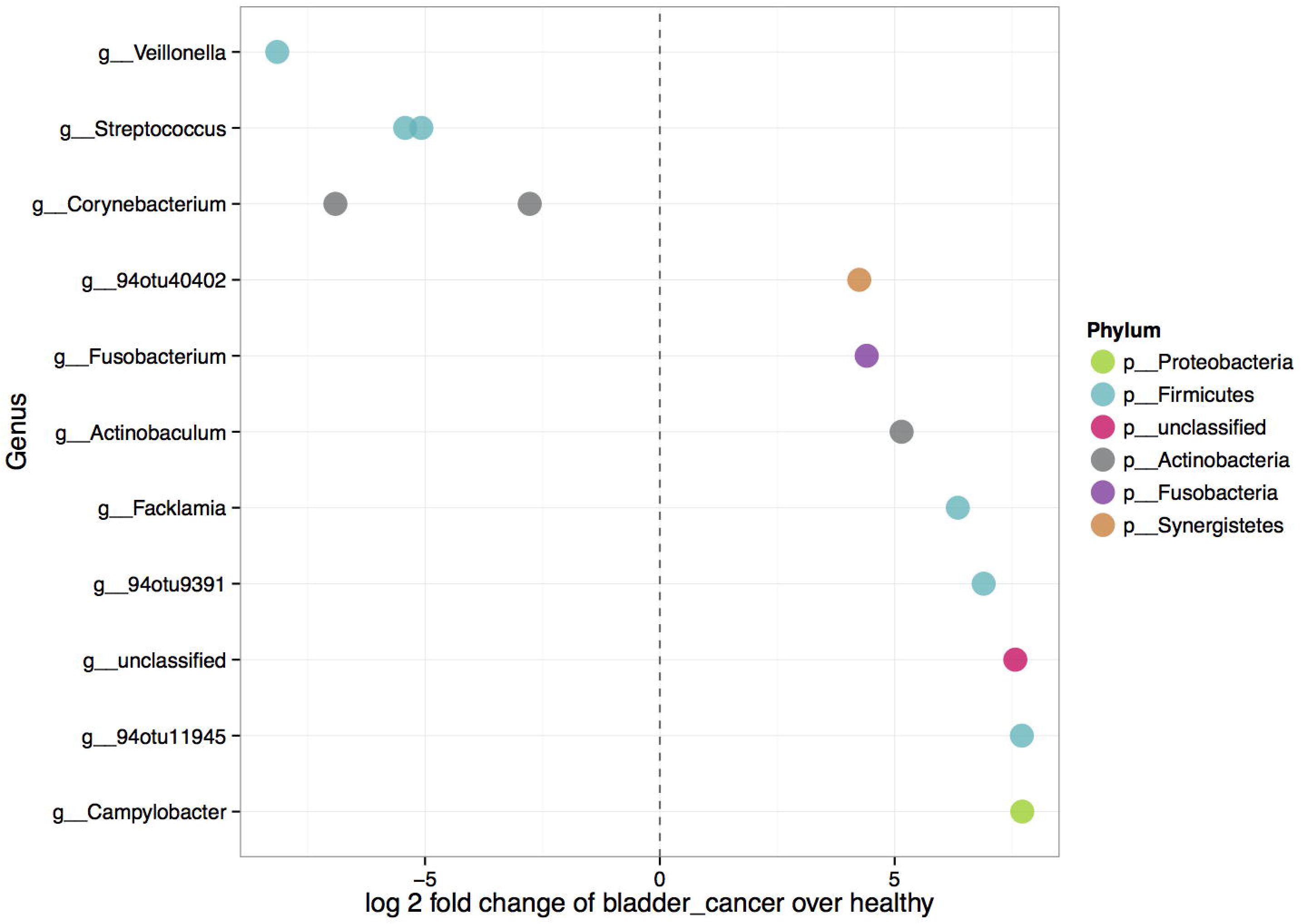
Differently abundant features between urine samples from bladder cancer patients and healthy controls. Each point represents an OTU belonging to respective genus. 94otu4042 was identified as *Jonquetella anthropi*, while 94otu9391 and 94otu11945 belong to family *Ruminococcaceae*. Features were considered significant if their false discovery rate-corrected p-value was less than or equal to 0.05, and the absolute value of the log2 fold change was greater than or equal to 1.

## Discussion

In this study, we have characterized urinary microbial communities of male patients diagnosed with primary or recurrent bladder cancer and compared it with those of disease free, age-matched controls. Although we did not observe major differences in overall microbiome profiles, we identified several OTUs that were significantly over-represented in bladder cancer or healthy subgroup. These differences suggest a possible role for urinary microbiome in bladder cancer pathogenesis that merits further evaluation.

A reduction in microbial diversity in urine of bladder cancer patients was not detected in our study (Fig. 1). While reduced diversity in the gut has been commonly linked with the state of disease^24^, changes in microbiome diversity have not been consistently associated with urinary tract disorders. Reduced bacterial diversity was observed in interstitial cystitis^17^, increased diversity was found in urgency urinary incontinence^15^ and chronic prostatitis^21^, while no changes in microbial diversity could be associated with overactive^12^ or neuropathic bladder symptomatology^18^. It may be that the abundance of specific bacteria in urine is more important than the total number of bacterial taxa present.

The most abundant phylum in both bladder cancer and healthy group was *Firmicutes*, followed by *Actinobacteria, Bacteroidetes* and *Proteobacteria* (Fig. 2), which is in good agreement with the composition of male urinary microbiome reported in current literature^10,20^. Also, inter-individual variability, observed between microbiomes of the participants in our study, has been repeatedly demonstrated in publications on urinary microbiome ^10,11,19,20^, which makes it challenging to define what constitutes a ‘core’ bladder microbiome.

Our comparison of bladder cancer versus healthy urine samples revealed bacterial taxa that were overrepresented in one of the sample subgroups (Fig. 4). Of note, among those OTUs that were more abundant in bladder cancer patients, an OTU belonging to genus *Fusobacterium* was detected. *Fusobacterium* is a common constituent of oral microbiome, but also an opportunistic pathogen with recognized carcinogenic potential^25,26^. It has been found associated with colorectal cancer in several studies^27,28,29,30^. Fusobacterial DNA was also detected in pancreatic^31^, breast^32^, esophageal^33^ and laryngeal^34^ cancer tissue, but this is to our knowledge, the first report on association of *Fusobacterium* with urothelial carcinomas. A proposed mechanism by which *Fusobacterium nucleatum* drives tumorigenesis involves activation of β-catenin cell signalling pathway leading to cell proliferation ^35,36^. A study by Kostić *et al*. ^37^, based on human sample analysis and mice model, suggests that, through recruitment of tumour-infiltrating immune cells, *Fusobacteria* generate a proinflammatory environment which supports tumour progression. There is also evidence that *F. nucleatum* inhibits NK cell cytotoxicity and T cell activity, thereby promoting immune evasion, which is one of the hallmarks of cancer^38^.

Enrichment of *F. nucleatum* in cancerous tissue seems to be mediated by binding of fusobacterial Fap2 lectin to tumour-displayed D-galactose-β(1-3)-N-acetyl-D-galactosamine (Gal-GalN Ac)^39^. A recently published pilot study demonstrated that in addition to colorectal carcinomas, various tumour types display Gal-GalNAc^40^. Although only moderate levels of Gal-GalNAc were found in urothelial carcinomas, the microbiome analysis undertaken in our study suggests that bladder cancer tissue can also be colonized by *F. nuclatum*.

An interesting observation on *F. nucleatum* is not to be overlooked: through action of its bacterial metabolites, *F. nucleatum* may be able to reactivate latent viral infections. In particular, *F. nucleatum* induced lytic replication of Kaposi’s sarcoma-associated herpesvirus^41^, which we found was associated with bladder cancer in our previous study^42^.

This indicates a possible synergistic interplay of bacterial and viral pathogens in bladder cancer development.

The most prominent OTU enriched in bladder cancer urines in our study (Fig. 4) was identified as *Campylobacter hominis*. Studies have shown that *Campylobacter* species are potentially pathogenic as they are able to produce toxins, invade epithelial cells, and avoid host immune responses. Similarly, *Campylobacter* species were found over-represented together with *Fusobacterium* in colorectal cancer^43^ and esophageal biopsies^44^.

Amongst other OTUs that were significantly more abundant in cancer patients, an OTU belonging to genera *Jonquetella* was detected. *Jonquetella* presence is potentially characteristic of urine microbiomes from individuals aged 70 and older^10^. Given the relatively small sample size, it is interesting that we could note clustering of samples according to age (Fig. 3), even though the study cohort already consisted of older individuals (Supplementary Table S1). These results confirm previous conclusion by Lewis at al. that aging modifies the composition of microbial communities in both the bladder and the gut^10^. Whether these shifts in microbiome composition that occur with aging increase cancer risks remains to be investigated.

Apart from being oncogenic, commensal microbiota may also provide beneficial, tumour-suppressive effects to the human host^7^. The concept that specific bacteria could protect against development of a malignant disease is particularly straightforward when considering urinary bladder cancer, because this is the only malignancy treated by a live microorganism, *Mycobacterium bovis* bacille Calmette-Guérin (BCG). Despite being used for more than 40 years, the molecular details of its therapeutic action are not fully elucidated. A proposed model suggests that BCG attaches to urothelial cells, which is followed by BCG internalization by bladder cancer cells and initiation of immune responses that destroy cancerous tissue^45^.

It could be envisioned that similarly to BCG, certain commensal bacteria, residing naturally in the healthy bladder, could serve the function of tumour surveillance or act beneficially in a different manner. In this study, five OTUs were found to be increased in healthy bladder and they included *Streptococcus, Veillonella*, and *Corynebacterium* species. *Streptococcus*, and to a lesser extent *Veillonella*, have repeatedly been observed in urine of healthy men^9,19,46^. In addition, characterization of microbial populations in specimens of another urological malignancy, prostate cancer, also showed the statistically significant enrichment of *Streptococcus* in nontumorous tissue^47^. *Corynebacterium* might be a typical urine component in healthy men as opposed to *Lactobacillus* which is prevalent in women^19^.

Our study on bladder cancer microbiome had some limitations. We used clean-catch midstream urine to sample bladder microbiome, which has its limitations but from an ethical point of view is a far less invasive method then bladder tissue biopsy or catheterization.

This study included only male patients. Men are at considerably higher risk of developing bladder cancer, which may be explained by the effects of sex hormones or gender differences in metabolic detoxification of carcinogens^48,49^. It would be interesting to explore if any specific member in the female urinary microbiome, which is dominated by *Lactobacillus* species, provides a protective effect.

As with other studies comparing disease versus healthy microbiome, it is not possible to say whether the microbial alterations are the cause or the consequence of the disease. Further longitudinal studies with a larger sample number at different stages of tumorigenesis and animal model studies will be needed to clarify the role of microbiome in bladder cancer formation and progression.

The main strength of the study is the novel insight on subtle changes of urinary microbiome in bladder cancer, including the increased abundance of possible pathogenic *Fusobacterium* species. Additionally, these results are important because the male urinary microbiome is often overlooked in urological studies, as the field is much more focused on female urogenital pathologies. More studies like this are needed to further define the core microbiome in both sexes and evaluate how it changes in specific disease states.

In conclusion, the 16S rDNA gene sequencing-based approach used in this work enabled us to characterize urinary bladder microbiome and detect differences in the relative abundance of specific bacteria in bladder cancer patients, with *Fusobacteria* as a possibly important representative. Whether observed differences contribute to bladder cancer development remains to be elucidated. A better understanding of the role of microbiome in bladder cancer could direct urologists to novel diagnostic and prognostic options, as well as to more personalized treatments and microbiome-targeted therapeutic interventions.

## Methods

### Subject recruitment and sample collection

The study began following approval from the Ethics Committee of the University Hospital in Split. Thirty six Caucasian men were recruited at the Department of Urology, University Hospital Split, between October 2015 and October 2016. The bladder cancer group contained 17 males diagnosed with primary or recurrent bladder cancer, and the control group had 19 healthy individuals who visited a urologist for prostate cancer screening check-up. None of the healthy controls had prostate cancer or indications for prostate biopsy (Supplementary Table S1). All experiments were performed in accordance with relevant guidelines and regulations and participants gave written informed consent for urine collection and analysis for research purposes. Exclusion criteria for both groups were antibiotic usage for at least one month prior to urine collection, positive history of sexually transmitted or recent urinary infections, diabetes and obesity. Additional exclusion criteria for bladder cancer patients were muscle-invasive disease, and previous treatment with Bacillus Calmette-Guérin (BCG) or radiotherapy. Tumour samples were evaluated by certified pathologist for malignancy grade and tumour stage. Clean catch, midstream urine was collected from all participants and stored at −80°C.

### DNA isolation from urine

Urine specimens (30 ml) were thawed and centrifuged at 7500 g, 4 °C for 10 minutes. The pellet was used for DNA extraction using PowerSoil@DNA Isolation Kit (MoBio Laboratories, Inc.), performed according to manufacturer’s protocol. To avoid environmental contamination, all isolations from urine samples and from the reagent-only extraction control were carried out within a PCR hood. Isolated DNA samples were placed at −20 °C until PCR amplification. DNA was quantified via the Qubit@ Quant-iT dsDNA High Sensitivity Kit 7 (Invitrogen, Life Technologies).

### 16S rRNA gene library preparation and MiSeq sequencing

PCR amplification of 16S rDNA, sequencing and analyses were performed by Second Genome. 16S rRNA gene V4 region was amplified with 515F-806R fusion primers that incorporate Illumina adapters and indexing barcodes^50^. PCR products were quantified using Quant-iT™ PicoGreen™ dsDNA Assay Kit from Invitrogen (Life Technologies, Grand Island, NY), pooled in equal molar ratios, and sequenced for 2 × 250 cycles on the Illumina MiSeq platform (Illumina, San Diego, CA).

### Bioinformatics and statistical analyses

Sequenced paired-end reads were processed using USEARCH^51^. All sequences hitting a unique strain in an in-house strains database with an identity ≥99% were assigned a strain Operational Taxonomic Unit (OTU). The remaining non-strain sequences were quality filtered, dereplicated and then clustered at 97% by UPARSE^52^. Representative OTU sequences were assigned a taxonomic classification at 80% confidence cutoff via mothur’s bayesian classifier^53^, against the Greengenes reference database of 16S rRNA gene sequences^54^ clustered at 99% OTUs. A prevalence filter was used to remove spurious OTUs that were observed in less than 10% of the sample set.

Diversity within samples (alpha diversity) was evaluated as richness and Shannon diversity. Richness is the number of observed unique Operational Taxonomic Units (OTUs), and Shannon Index evaluates richness and the abundance of each OTU (evenness). Dissimilarity between samples (beta diversity) was assessed using the Bray-Curtis dissimilarity measure ^55^. To visualize inter-sample relationships, Principal Coordinates Analysis (PCoA) was performed.

Differences in the overall microbial composition between bladder cancer and healthy samples were assessed by permutational analysis of variance, PERMANOVA^56^. To identify taxa that were significantly different between bladder cancer and healthy samples, we used DESeq2 package^57^, described for microbiome applications^58^. DESeq2 was run under default settings and q-values were calculated with the Benjamini-Hochberg procedure to correct p-values and control for false discovery rates.

## Data Availability

The datasets generated during the current study are available in the European Nucleotide Archive, accession number: PRJEB22327

## Acknowledgements

This work was supported by Croatian Science Foundation Grant No: IP-2014-09-1904 to JT. The authors thank to Ms. Sandra Vujević for technical assistance.

## Author Contributions

JT, VBP and MŠ designed the study, MŠ enrolled the participants and collected samples, VBP, CTC, LSC and BR collected the data, CTC and LSC analysed the sequencing data, VBP drafted the main manuscript text, CTC and LSC prepared the figures and all authors contributed to the final article.

